# PROLONG: Penalized Regression for Outcome guided Longitudinal Omics analysis with Network and Group constraints

**DOI:** 10.1101/2023.11.06.565845

**Authors:** Steve Broll, Sumanta Basu, Myung Hee Lee, Martin T. Wells

## Abstract

**Motivation:** There is a growing interest in longitudinal omics data, but there are gaps in existing methodology in the high-dimensional setting. This paper focuses on selecting metabolites that co-vary with Tuberculosis mycobacterial load. The proposed method is applied to general continuous longitudinal outcomes with continuous longitudinal omics predictors. Simple longitudinal models examining a single omic predictor at a time do not leverage the correlation across predictors, thus losing power. We propose a penalized regression approach on the first differences of the data that extends the lasso + Laplacian method (Li and Li 2008) to a longitudinal group lasso + Laplacian approach. Our method, PROLONG, leverages the first differences of the data to address the piecewise linear structure and the observed time dependence. The Laplacian network constraint incorporates the dependence structure of the predictors, and the group lasso constraint induces sparsity while grouping metabolites across their first differenced observations.

**Results:** With an automated selection of model hyper-parameters, PROLONG correctly selects target metabolites with high specificity and sensitivity across simulation scenarios and sizes. PROLONG selects a set of metabolites from the real data that includes interesting targets identified during EDA.

**Availability:** R package ‘prolong’ is in development.

**Conclusions:** PROLONG is a powerful method for selecting interesting features in high dimensional longitudinal omics data that co-vary with some continuous clinical outcome.

## 1 Introduction

An emerging problem in omics research is the discovery of useful biomarkers for repeated-measure data where both the clinical outcome and a large number of omics predictors are measured a few times on a number of individuals. In many clinical studies of disease progression, prognosis, and effectiveness of new treatments, we track clinical outcomes of a small group of patients over a handful of times (a few weeks, *t* = 3 *−* 8). Recently it is also more feasible to collect data on a large number of omics features (metabolites, gene expressions, etc.). Clinicians with access to this large pool of omics features are interested in building computational biomarkers predictive of the clinical outcome because these data can be useful both as non-invasive markers of the disease process and as insight into biological pathways [1] [2] [3].

Our work is motivated by a pulmonary tuberculosis study consisting of 4 time point observations for 15 subjects each with measured sputum mycobacterial load, quantified by the time in hours to Mtb culture positivity (TTP), and abundances of up to 915 urinary metabolites. Previous work has been performed on the identification of urinary metabolite-based biomarkers for patients with pulmonary tuberculosis [4] [5] and creating a general methodology for metabolite-based biomarker discovery in settings with longitudinal continuous outcomes.

Existing works in the clinical omics literature use univariate longitudinal predictive regression models [6] and [7] for the clinical outcome of interest. Typically, variable selection is conducted by comparing the p-values of the univariate models. Beyond the usual multiplicity adjustment limitations of one-at-a-time modeling and testing, we argue that, by failing to incorporate the underlying omic feature and time correlation structures, this approach is susceptible to low sensitivity and statistical power.

The literature on high-dimensional omics in molecular biology and genomics is vast. However, these methods are not designed around using a time-varying clinical outcome to guide variable selection. Existing highdimensional omics approaches, whether for single or repeated measures, generally focus on differential expression between two or more treatment groups (DESeq2 [8], limma [9], OmicsLonDA [10]). Additional challenges arise in our data that are not always sufficiently addressed in the high-dimensional omics setting. The data has low signal but high variability across a few subjects and across 4 time points. Auto-correlation is evident in the data and must also be accounted for.

To fill this gap we propose the PROLONG method, which can jointly select longitudinal features that co-vary with a time-varying outcome on the first-difference scale. Network and group lasso constraints induce sparsity while also grouping the different time components of each metabolite and incorporating the correlation structure of the data. Working in the first-difference scale controls for the temporal dependence inherent in the data, improving power and avoiding time-related assumptions.

The incorporation of a outcome, rather than just identifying molecules whose abundance differs between treatment groups, allows PROLONG to identify molecules that either co-vary with the outcome or are significantly correlated on the first-differenced scale with those that do. This allows us to focus on successful treatments and identify a list of target molecules that may be mechanistically involved in improvement in outcome.

PROLONG performs very well in simulations with both high sensitivity and specificity with both uncorrelated and correlated simulated data. PROLONG is also tested on the data of the motivating example of the metabolite abundances as potential predictors and mycobacterial load as our clinical outcome, shown in Figure 1. For this data, PROLONG is stable across both parameters and selects a set of target metabolites identified in exploratory data analysis and by clinicians, while the univariate longitudinal model picks out no metabolites at reasonable FDR thresholds. PROLONG achieves this performance despite having only 15 subjects for 352 metabolite measurements replicated 4 times each.

**Figure 1:**
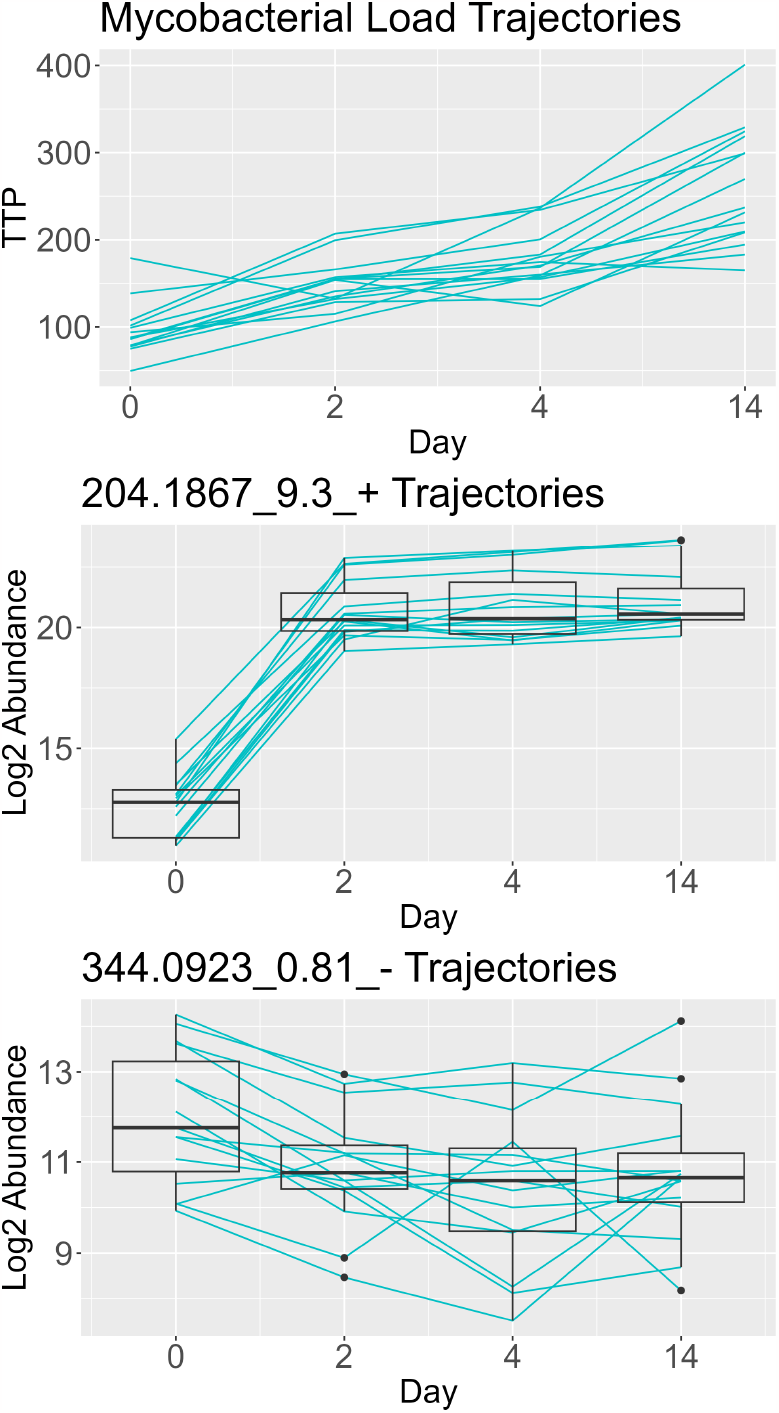
Trajectories for Time to Positivity which is inversely related to mycobacterial load (top), an interesting metabolite (middle), and a more stationary but still potentially interesting metabolite (bottom).

Some of the metabolites selected by PROLONG were identified as targets from the exploratory analysis phase due to their sharp change from the baseline to the second measurement, like 204.1867 9.3+. whose trajectories are shown in Figure 1. However, most metabolite trajectories are not cleanly separated between starting and final measurements as for 204.1867 9.3+. Many metabolites do not change significantly in mean over time, such as 344.0923 0.81 shown in Figure 1. These metabolites have different levels of variation across time points but do not show a sustained long-term effect after treatment. Altogether, these characteristics of the data prove challenging for univariate mixed effects models and make a high-dimensional method like PROLONG a necessity.

## 2 Methods

To provide motivation for PROLONG, we first describe the univariate mixed effects model often used for longitudinal omics regression [6]. Then we provide our proposed univariate alternative model and a natural hypothesis testing technique. We end this section with our proposed multivariate extension, which includes constraints to induce sparsity in the high-dimension scenario and to incorporate the dependence structure of the omics predictors.

### 2.1 Background: mixed effects models

This article’s motivating example is a short-term pulmonary tuberculosis drug trial. The prototypical data from such a short-term trial consists of regressors 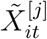 and response 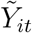 for subject *i* = 1, …, *n*, omics feature *j* = 1, …, *p*, and time point *t* = 1, …, *T*. A univariate linear mixed effects model for the levels of 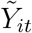, with fixed effects for the omics feature and for the time component and a subject-specific fixed or random effect, then takes the form

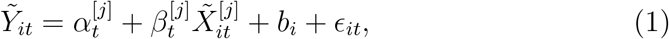

where 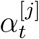 is our mean response across subjects at time *t, b*_*i*_ is the subject specific fixed or random effect, *ϵ*_*it*_ is the mean-zero error term and 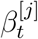 is the time-dependent omics feature-specific parameter of interest.

Fixed effects are generally preferred for levels modelling when understanding how individual entities change over time while controlling for time-invariant heterogeneity. Random effect models are appropriate when the focus is on estimating population-level parameters and the main interest lies in the variation across entities. However, if the individual-specific effects significantly correlate with the explanatory variables, using random effects can lead to biased and inconsistent estimates. The choice between fixed and random effects models depends on the presence and nature of unobserved heterogeneity in the panel data. If one suspects unobserved factors correlate with the explanatory variables, a fixed effects model may be more appropriate to control for this correlation. On the other hand, if you believe that unobserved factors are uncorrelated with the explanatory variables, a random effects model might be more efficient and suitable.

Longitudinal fixed and random effects models account for potential temporal dependence in the data. Still, they may have insufficient power to select true metabolites with short-term changes and trends over time in data settings. With the TTP trajectories, there are some variance changes over time across subjects, but most of the ‘signal’ is the consistent, substantial increase at every time point. This is particularly relevant when the time course data is primarily focused on short-term dynamics, and controlling for time-invariant unobservable factors is crucial [11].

To avoid having to work around temporal dependence and to try to emphasize the variation in the change from time point to time point in the levels model, we move to a first-difference, or delta scale, in the 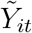. When we regress the first differences of the outcome on our first differences of the metabolites, we seek to determine which metabolites change over time across subjects in a way that linearly corresponds to how the outcome measure changes over time across subjects. This first differences approach should be used when there are concerns about unobserved time-invariant heterogeneity and researchers want to focus on short-term changes and trends over time. This is particularly relevant when the panel data exhibit high levels of time-persistent unobservable characteristics that might bias the estimates in the fixed effects model. The first difference approach also effectively deals with potential endogeneity concerns related to time-invariant unobserved factors. The first-difference approach is appropriate for our motivating example, as the subjects all receive the same treatment but show time heterogeneous TPP and metabolite abundances. We view our independent variables not as a treatment themselves but as potential mediators for the treatment effect in each subject. Identifying significant linear relationships on the delta scale can inform clinicians of biomarkers that may be involved in improving outcomes from the treatment.

### 2.2 PROLONG: Univariate Case

In this subsection we give the sequence of step to deduce a matrix representation of the differenced regression model from (1). For each subject, we first matricize the data to incorporate all time points into a single model. Once in the form of matrices, the difference between subsequent time points can be represented in terms of other matrices.

Given our original model in (1), the *n × T* response matrix 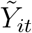 can be represented and differenced in the following step

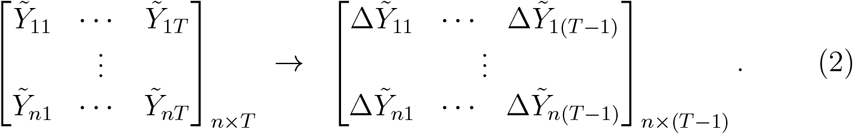

where each 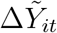 is the first-difference 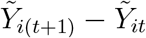.

Vectorizing this matrix yields the *n*(*T −* 1) *×* 1 vector *Y* to be used as the dependent variable in our differenced model where

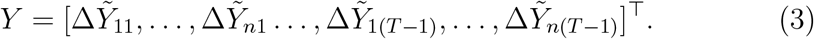

For the *j*^th^ metabolite (*j* = 1, …, *p*), the *n × T* matrix can be represented and differenced in the following step

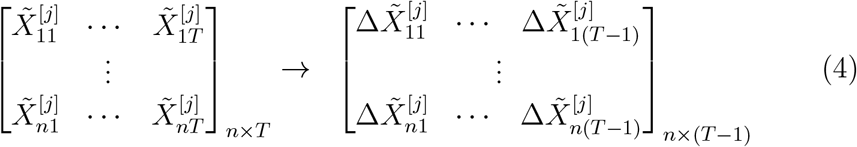

where each 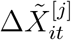 is again the first-difference 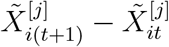.

To derive the difference design matrix we need to respect the temporality of the observations and regress 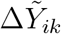 for *k* = 1, …, *T −* 1 on the differenced measurements of 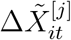 for *t* = 1, …, *k* and *j* = 1, …, *p*. Consequently for the *j*^th^ metabolite the *n*(*T −* 1) *× T* (*T −* 1)*/*2 design matrix is

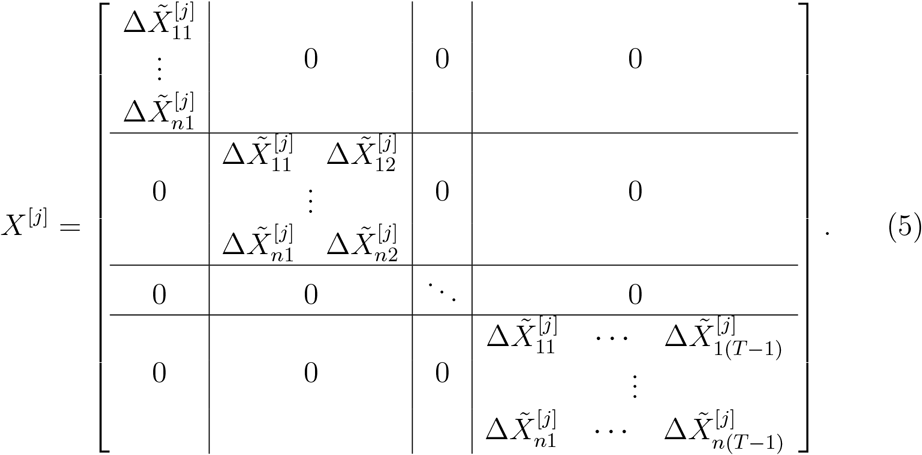

An appropriate univariate test for whether the effect of a metabolite on the outcome *Y* is the Wald test [12] for the coefficients from the model

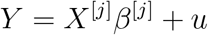

where *u* is the mean-zero error vector, and

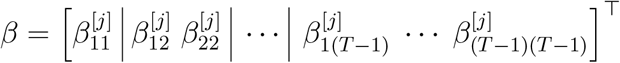

For the test parameters we use the *T* (*T−* 1)*/*2 coefficients from the (*T −*1) models for each component of *Y*, and for the covariance matrix we use a block diagonal with (*T −*1) blocks containing the covariance matrix for the coefficients of the respective models. This univariate test encapsulates the model structure we are proposing and could be used for small sets of variables that are not heavily correlated. For larger or correlated variable sets we will need a joint model with appropriate constraints.

Note that the design matrix in (5) does not have a column of the ones that would represent an intercept. When the intercept in the levels model (1) is a constant, say *α*_0_, the first-difference operators in (2) and (4) eliminates it. If one were to include a column of ones in the design matrix (5), one is essentially assuming *α*_*t*_ in the levels model (1) is varying linearly in *t*. If one were to assume that *α*_*t*_ in the levels model (1) contains a *k*^th^ order polynomial time trend, one could apply the (*k−* 1)^th^-order difference operator in (2) and (4) rather than first-order operators.

### 2.3 PROLONG: Multivariate Case

The above construction gives us the desired regression structure, though we now need to incorporate all metabolites simultaneously for a joint model. The easiest way to do so given our eventual construction of a correlation graph is to replace 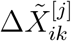 in *X*^[*j*]^ above with row vector

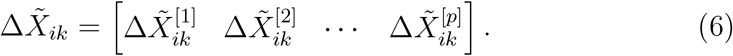

With these substitutions we get the differenced *n*(*T −*1) *×pT* (*T −*1)*/*2 design matrix

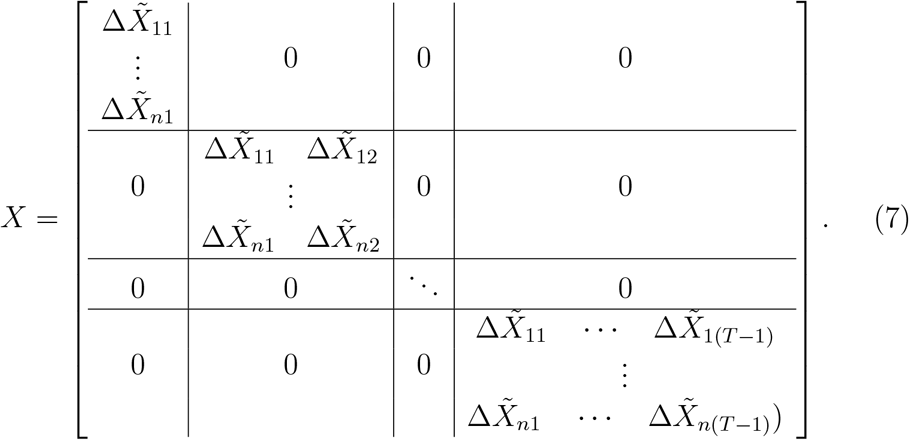

Using the matrix representations of the difference based dependent and independent variables in (3) and (7) the longitudinal regression model takes the form

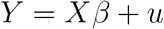

where *u* is the mean-zero error vector and

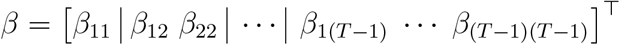

with 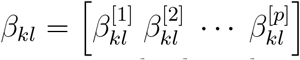 for 1 *≤ k ≤ l ≤ T −* 1

Here, we scale the columns of *X* such that for every column *j*. Note that this is a zero intercept model and that we do not center the columns of Δ*X*. The basic regression model does not contain a known underlying graph connecting the dependence among the independent variables (metabolites), so we now develop a graph-based penalization via the design matrix *X* dependence structure. Since each of the *T −* 1 separate time components of *Y* are jointly modeled the block diagonal dependence matrix will mirror our design matrix.

For *i*^th^ subject, define Δ*X*_*i*_ as the (*T −* 1) *× p* matrix with the *k*^th^ (*k* = 1, …, *T −* 1) row equal to (6). Note that Δ*X*_*i*_ = *vec*(*χ*_*i*_), where *χ*_*i*_ is (*T −* 1)*×p*. Dropping the *i* subscript, define a dependence matrix (such as Pearson or Spearman correlation, or Kendall’s *τ*) of *χ* to be

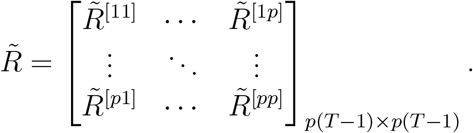

Each block 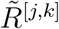 is a (*T −* 1) *×* (*T −* 1) dependence matrix for the columns of the *χ*, that is, the block of dependencies among the independent variables. Now consider the dependence matrix *ℛ* of *vec*(*χ*^*⊤*^). This is also a *p*(*T −* 1) *× p*(*T −* 1) matrix but can be written with blocks corresponding to pairs of time points of *χ*

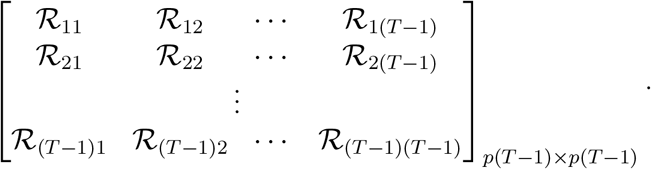

The dependence matrix associated with the design matrix *X* in (7) can be constructed using the blocks of *ℛ* as

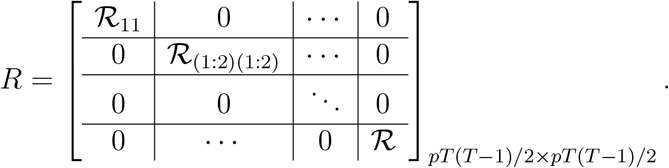

PROLONG uses a regular group lasso penalty along with a network constraint via the Laplacian matrix of the graph associated with *R*. First, estimate *R* via 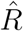. We define this graph *G* as having edges *e* = (*u ∼ v*) between columns *u, v* of *X* whose edge-weights are 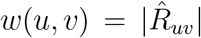. The degree of each vertex is

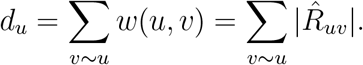

As in [13], we define the normalized Laplacian matrix *L* for graph *G* element-wise as

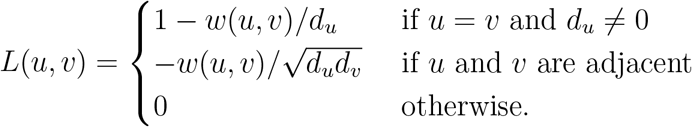

The penalization criterion used by [13] contains terms for the network constraint and for the lasso penalty, and for our data would be *λ*_1_ *β* _1_ + *λ*_2_*β*^*T*^ *Lβ*.

Following Lemma 1 in [13], given non-negative *λ*_1_, *λ*_2_, we create a new augmented dataset by appending a 0-vector to *Y* and *𝒮* ^*T*^ to *X*, where *𝒮* = G*D*^1*/*2^ for *L* = G*D*G^*T*^, resulting in (*𝒴, 𝒳*)

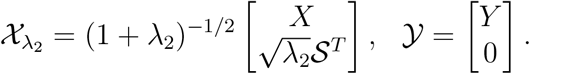

Operationally, we estimate the vector *β*^*\**^ = *β*(1 + *λ*_2_)^1*/*2^ via lasso and then adjust by the factor (1 + *λ*_2_)^*−*1*/*2^ to debias the augmented model.

The group lasso penalty [14] is defined as 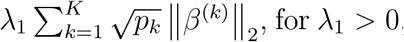, for *λ*_1_ *>* 0, where 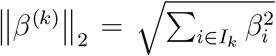, for each *k*, the index for group and *p*_*k*_ is the size of group *k*. Put this all together, the group lasso + Laplacian penalty is

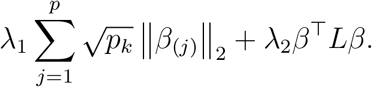

As in the lasso + Laplacian model, compute *𝒮* from the G*D*G^*T*^ decomposition of *L* and construct augmented dataset 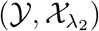. We apply the group lasso penalty using group-wise-majorization-descent (GMD) [15] implemented in the R package ‘gglasso’ [16].

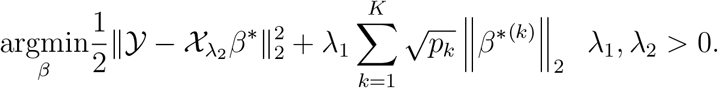

Again upon adjusting for augmented model representation, the obtained solution is multiplies by (1 + *λ*_2_)^*−*1*/*2^.

The tuning parameter *λ*_2_ is selected via MLE, using the following optimization problem from [17], that is, *λ*_2_ is the solution of the maximization of 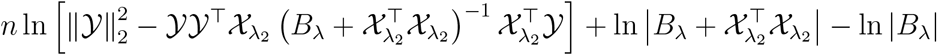, where *B*_*λ*_ = *λ*_2_*L* + *λ*_*R*_*I*. The *λ*_*R*_*I* term ensures *L* becomes invertible which is needed for the MLE derivation.

After minimizing over both *λ*_2_ and *λ*_*R*_, we add *λ*_*R*_*I* to *L* before computing *𝒮* via G*D*G^*T*^ decomposition. Then *λ*_1_ is selected via group lasso crossvalidation on the augmented model (𝒴, 𝒳) generated using *λ*_2_ and *λ*_*R*_.

PROLONG inherits characteristics of both network-constrained regularization and group-lasso. Most importantly, the network constraint guarantees that if *x*_*i*_ = *x*_*j*_ then 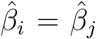, and omics features that are highly correlated but not identical will be grouped together by the constraint. More details and asymptotic properties are discussed by [13]. The group penalty ensures that all coefficients associated with a given omics feature are either zero or non-zero, but not necessarily identical. Comparing the coefficients within a selected group can provide insight into the relative predictive strength of the feature at different measurement times for the feature and for the outcome. Grouping together all 𝒳columns associated with a given omics feature provides clarity in determining which omics features are ‘selected’ by the model.

## 3. Simulation Scenarios and Results

First, we will compare the performance of PROLONG to that of our benchmark univariate mixed effects model for simulated uncorrelated data. Uncorrelated data is a good first comparison as the network constraint portion of PROLONG will not have any real signal to work with, so it could be thought of as more challenging than data with correlated predictors.

We are not so much interested in recovering the exact generating coefficients as we are in correctly distinguishing between targets and noise. To assess the performance of PROLONG and the univariate method we will compare the selection rates for each variable, target and noise, across 100 simulations.

### 3.1 Fully Simulated Data with Uncorrelated Predictors

The following simulation setup emulates aspects of the real data with varying levels of signal. First, we show results for a small-scale scenario with 10 targets and 20 noise variables, then for a larger-scale scenario with 20 targets and 80 noise variables using the same data generating scheme. The data-generating scheme sets *β* = (1*/*3, …, 1*/*3, 0, …, 0) with

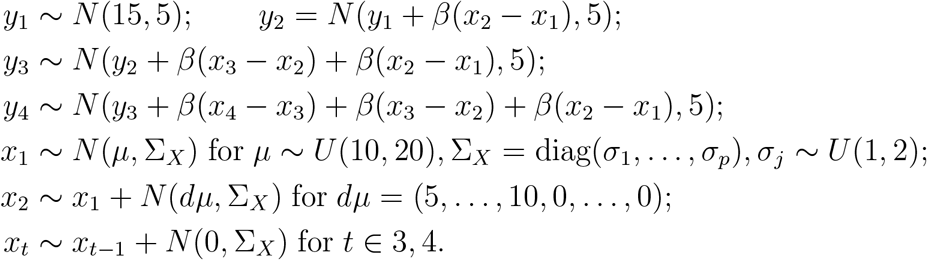

#### 3.1.1 Small Scale Results

As seen in figure 2(a), with 10 uncorrelated target variables and 20 noise variables, we find that the univariate mixed effects models struggled to distinguish between the targets and noise. This pattern holds across FDR thresholds, as the rate of acceptance increased flatly across targets and noise. PROLONG picked out every target in 98 simulations and missed 1 target each in the other 2, and selected 7.4% of the noise across simulations. At an FDR level of 0.05, the Wald test selected every target across all simulations and picked up 11.45% of the noise across simulations. At an FDR of 0.01, Wald tests picked up only 6.6% of the noise, slightly outperforming PROLONG on this small-scale data.

**Figure 2:**
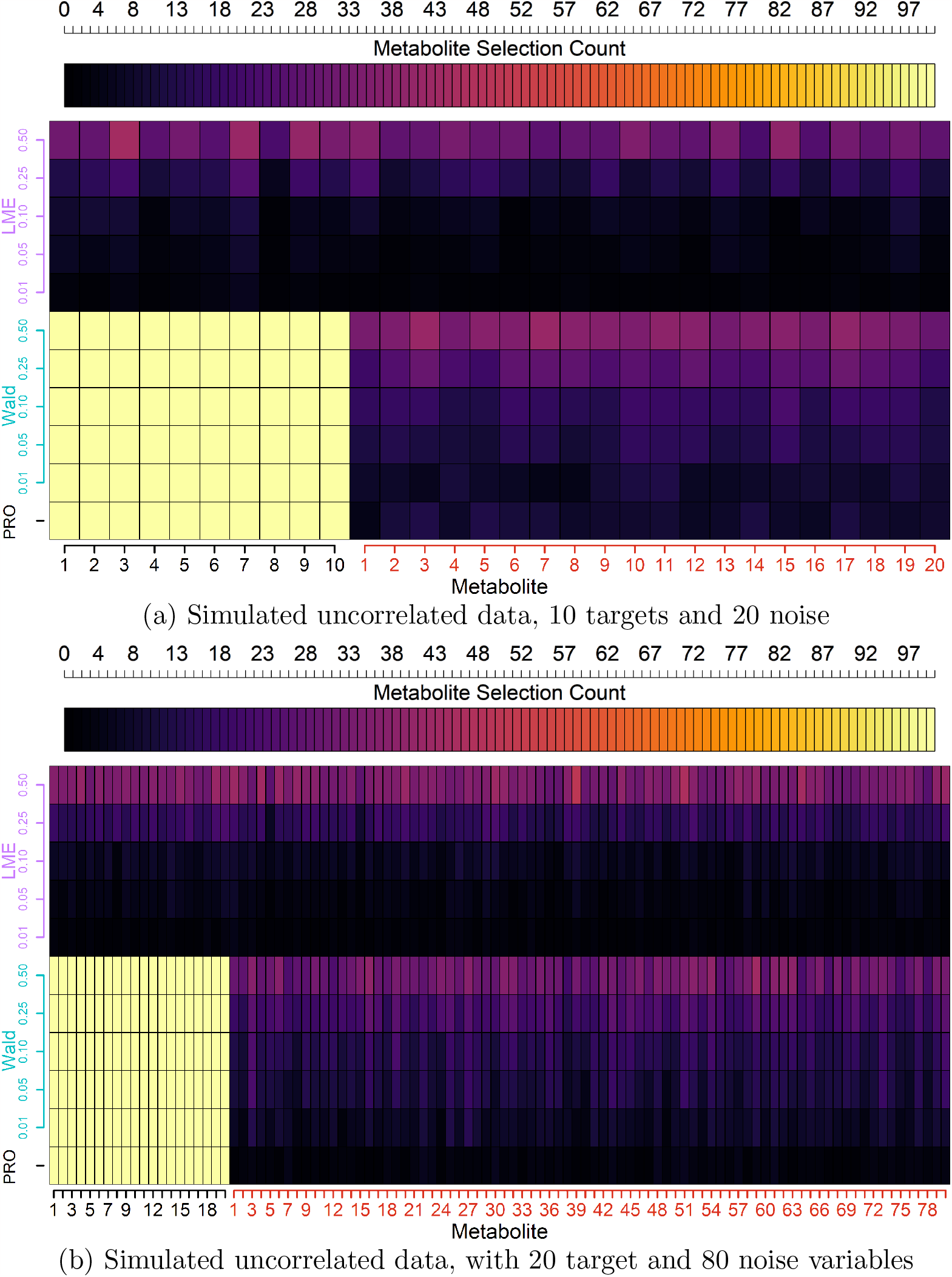
Comparison of selection counts (top axis) across simulations for target and noise metabolites (x-axis) using PROLONG, univariate Wald tests, and univariate Linear Mixed Effects (LME) models at Various FDR Thresholds (y-axis).

Figure 3 shows the coefficient paths and group lasso CV loss for a simulation run from each of the correlated and uncorrelated small-scale simulations. The blue lines show the coefficient paths for the target variables, red show the paths for the noise variables. The higher blue paths indicate coefficients for the time points that are used to generate the changes in *Y*, while those closer to zero represent time points not used to generate *Y* but that are kept from being shrunk to zero by the group penalty. In this example of the uncorrelated data, we see a wide gap in the log lambda scale for the selection of noise and targets. We can observe a steady decline in the noise coefficients, while the non-significant time points for the target variables are relatively flat across the lambda scale.

**Figure 3:**
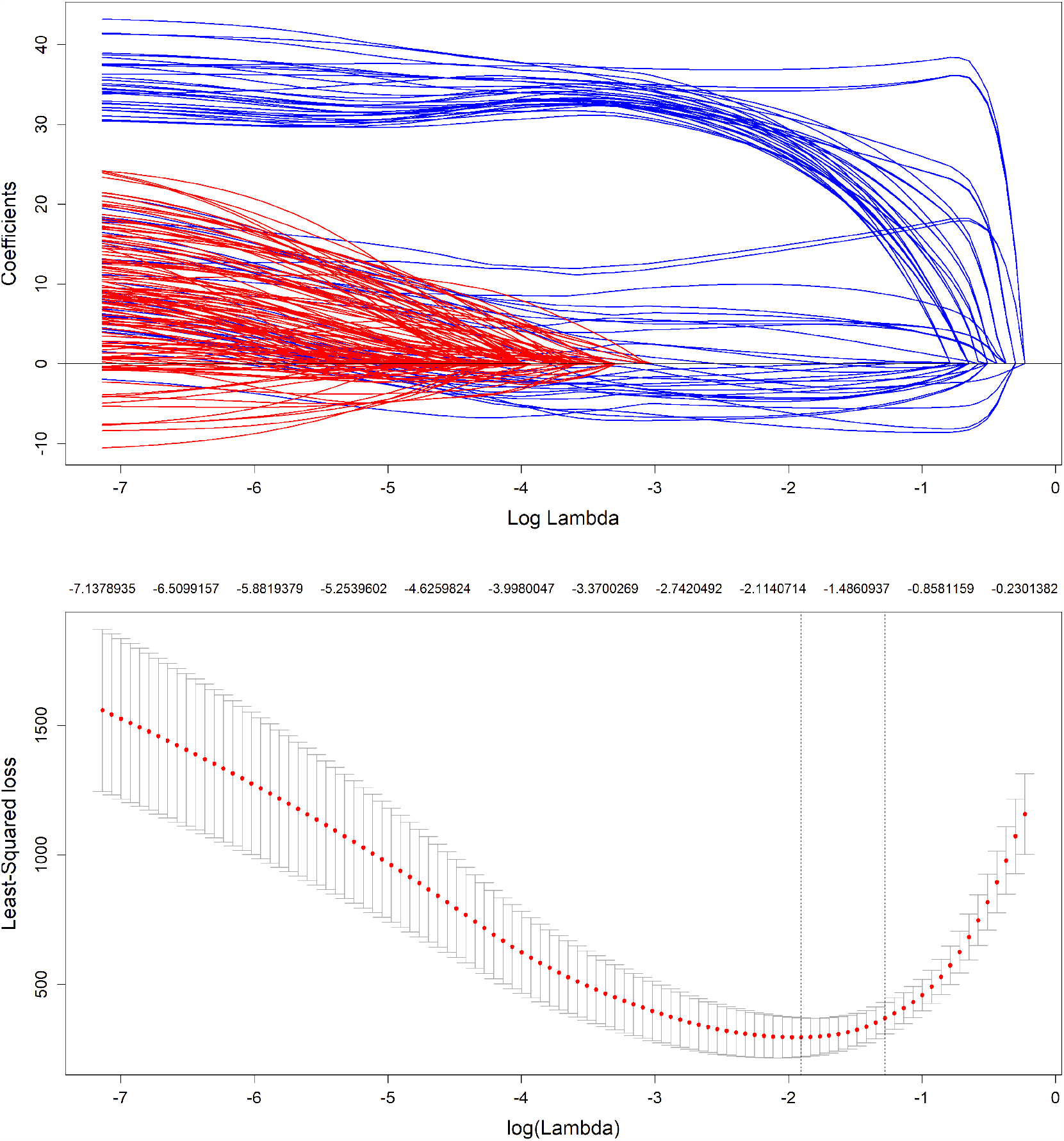
Top image shows coefficient trajectories across group lasso *λ*_1_ values for both target (blue) and noise (red) simulated metabolites. The bottom image shows LS loss as a function of the group lasso *λ*_1_

#### 3.1.2 Large Scale Results

Figure 4(b) shows that PROLONG selected every target metabolite but only 1.29% of the noise. The Wald tests also selected the targets perfectly at FDR levels of both 0.05 and 0.01 as well as 9.921% and 5.65% of the noise, respectively. With this larger variable set we see a much stronger performance for PROLONG compared to the Wald test. Again, the univariate mixed effects models struggled as expected with flat selection across noise and target variables.

**Figure 4:**
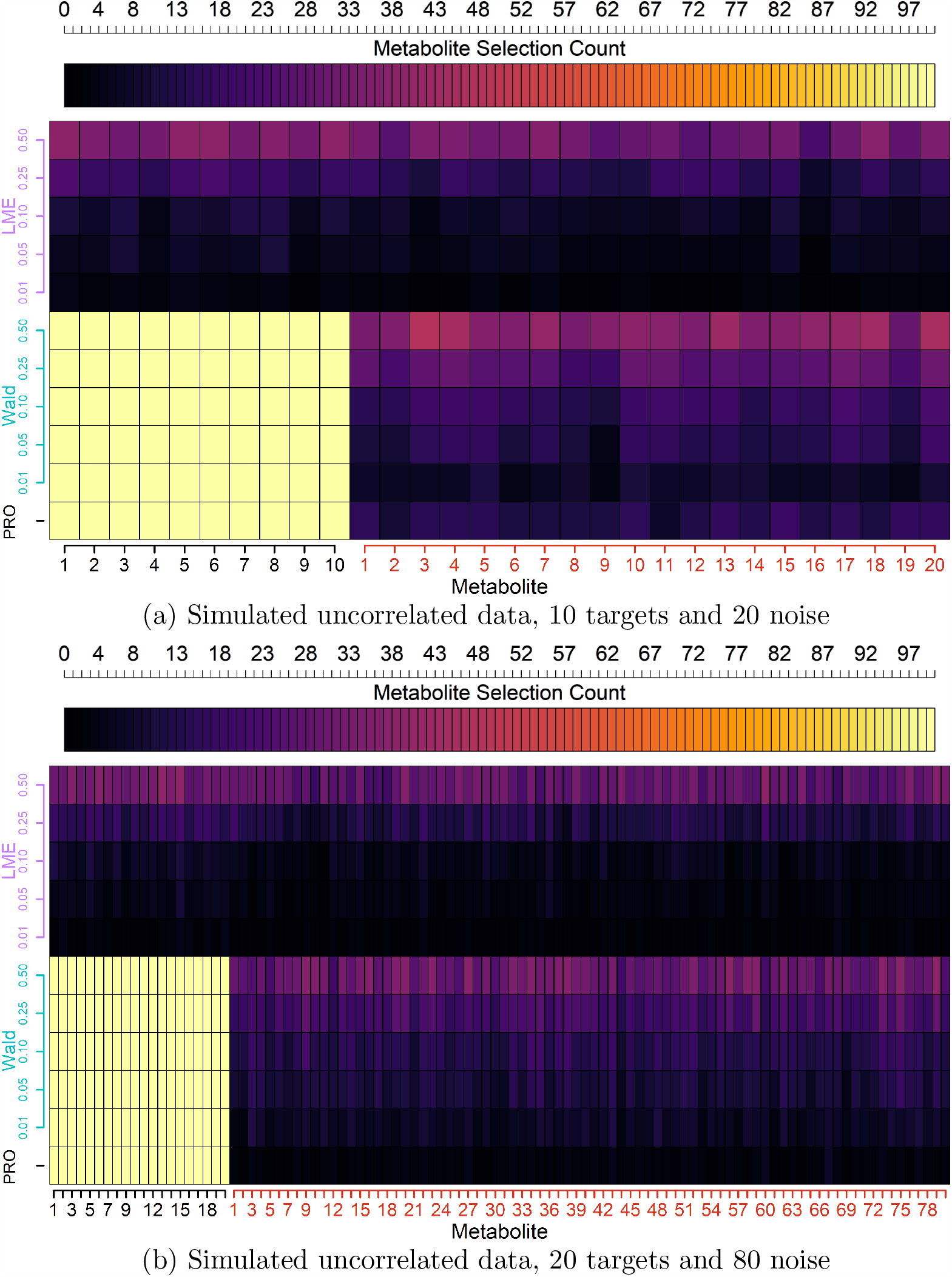
Comparison of selection counts (top axis) across simulations for target and noise metabolites (x-axis) using PROLONG, univariate Wald tests, and univariate Linear Mixed Effects (LME) models at Various FDR Thresholds (y-axis).

### 3.2 Fully Simulated Data with Correlated Predictors

Next, we will compare PROLONG and the univariate tests for performance on simulated correlated data. The univariate mixed effects models cannot make use of any correlation between variables, so we do not expect the results to vary meaningfully between uncorrelated and correlated simulations.

This simulation setup only differs from the previous by the definition of Σ_*X*_. Here we have

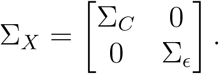

where for Σ_*d*_ = diag(*σ*_1_, …*σ*_*p*_) our covariance matrix Σ_*C*_ = Σ_*d*_*C*Σ_*d*_ associated with our target variables is constructed so that the variances on the diagonal are in the same range as those of the noise variables, while the off-diagonal elements correspond to off-diagonal elements of correlation matrix *C* drawn uniformly from (*−*1, 1).

Constructing Σ_*X*_ in this way ensures we have some (random) correlation between the target variables that generate *Y* that can be leveraged by the correlation network penalty.

#### 3.2.1 Small Scale Results

Figure 4(a) displays our results for 10 correlated target variables and 20 uncorrelated noise. The univariate mixed effects model struggled as in the uncorrelated case, while PROLONG and Wald tests were able to pick up every target across FDR thresholds. PROLONG selected 12.05% pf noise, slightly more than Wald FDR 0.05 at 11.95% and significantly more than Wald FDR 0.01 with 5.9% of noise.

#### 3.2.2 Large Scale Results

Figure 4(b) shows results for 20 correlated target variables and 80 uncorrelated noise variables. PROLONG, Wald FDR 0.05, and Wald FDR 0.01 selected all of the targets along with 1.21%, 5.43%, and 9.31% of noise respectively. Once again PROLONG beat out Wald with the larger dataset, and did slightly better than in the uncorrelated case.

#### 3.2.3 Full Data Scale Results

Figure 5 shows results for 50 correlated target variables and 300 uncorrelated noise variables. The univariate mixed effects models again showed no difference across target and noise. PROLONG, Wald FDR 0.05. and Wald FDR 0.01 select all targets once again, with 0.31%, 5.04%, and 8.6% of noise picked up by each respective model.

**Figure 5:**
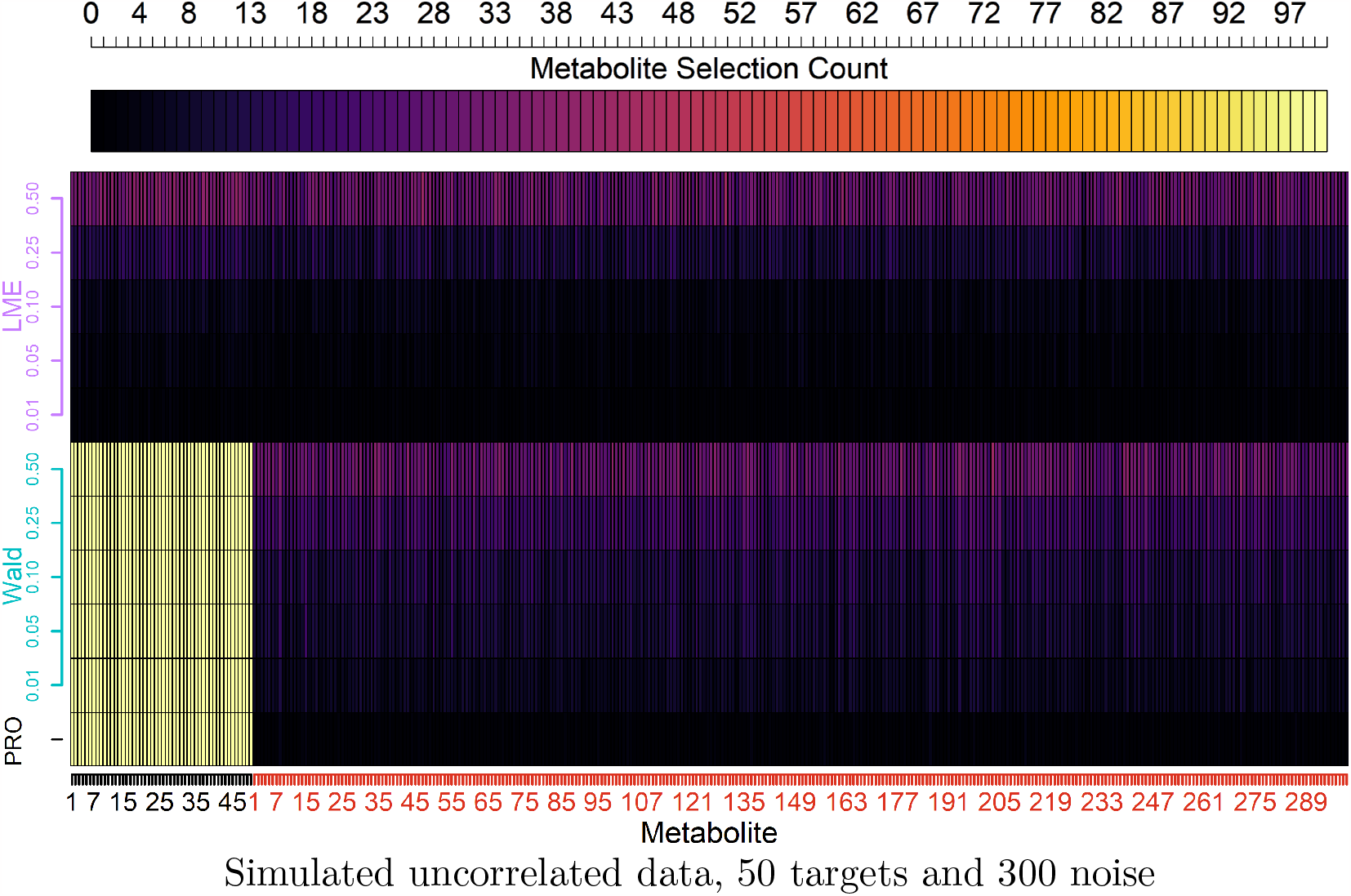
Comparison of selection counts (top axis) across simulations for target and noise metabolites (x-axis) using PROLONG, univariate Wald tests, and univariate Linear Mixed Effects (LME) models at Various FDR Thresholds (y-axis).

### 3.3 Summary of Results

Altogether, we can observe that PROLONG had near-perfect sensitivity in both correlated and uncorrelated simulated data and only improved as we increased our variable size and sparsity. The univariate mixed effects models struggled in all scenarios. The Wald tests at an FDR threshold of 0.05 performed similarly to PROLONG in the small-scale simulations, but only slightly decreased noise selection rate with increasing dimension while PROLONG drastically improved.

The correlation did not provide a significant advantage here, likely due to the strong signal relative to the average correlation strength. Even uncorrelated variables with a lower signal were picked up nearly always across all scenarios, so there was little room for improvement with the incorporation of a correlation structure within the target variables.

## 4 Data

### 4.1 Clinical Significance

An estimated 10 million people develop tuberculosis yearly, and 1.4 million die of pulmonary tuberculosis [18]. Since curing pulmonary tuberculosis is not always achieved in longer-term treatment regimes, reducing treatment to shorter regimes may improve adherence and increase treatment completion and cure rates [19].

Importantly, this TB dataset is not particularly special or unique. In fact, this data is reflective of an increasingly common collection of biomarkers across a few time points with some continuous outcome of interest. There is extensive work on differential expression methods, but here we have monotonically increasing but variable improvement in TTP across subjects, as well as potentially interesting biomarkers that are not significantly different from the first and last time points of measurement. While some metabolites do show a strong spike and plateau pattern, others primarily show variability between subjects at the first few time points while having a relatively stable mean. This data is representative of other new datasets while also providing a challenge that existing methods are not intended to address.

### 4.2 Collection and Preprocessing

Missingness is a common issue in mass-spectometry based metabolomics data [20], and so for the purpose of model development we limited the following analysis to metabolites with less than 20% observations missing across subjects and time points. We have 352 metabolites that meet this criterion, of which 79 have no missing values. We split the mostly complete 352 metabolites into 4 matrices, one for each time point, and used softImpute [21] [22] to fill in missing values for each time point’s matrix of metabolite abundances by subject. Imputation was done in this way to avoid introducing any new dependence across time points.

### 4.3 Benchmark Method

Mean squared error and similar model evaluation criteria are not particularly meaningful for this group lasso + Laplacian approach, so we instead investigate the selected metabolites for PROLONG and for univariate longitudinal mixed effects models, checking for target metabolites identified in our exploratory analysis and previous literature.

### 4.4 Results

No metabolites are selected by the univariate mixed effects models. This approach likely struggles with the apparent differential nature of the data, as the time intercepts tend to have much higher test statistics than the metabolite abundance terms.

Figure 6 shows 5 of the 45 metabolites selected by the PROLONG model, sorted by average coefficient magnitude but excluding 204.1867 9.3 + shown in the introduction. PROLONG also picks up 344.0923 0.01 shown in the 1. 100.1019 11.01 + shows a pattern similar to 204.1867 9.3 +, but among the rest we don’t always have the same clear spike and plateau. There are still easily observable differential effects in at least a few subjects that were picked up by the model. We can also observe that PROLONG does not only pick up metabolites that are differentially expressed between the first and last time points.

**Figure 6:**
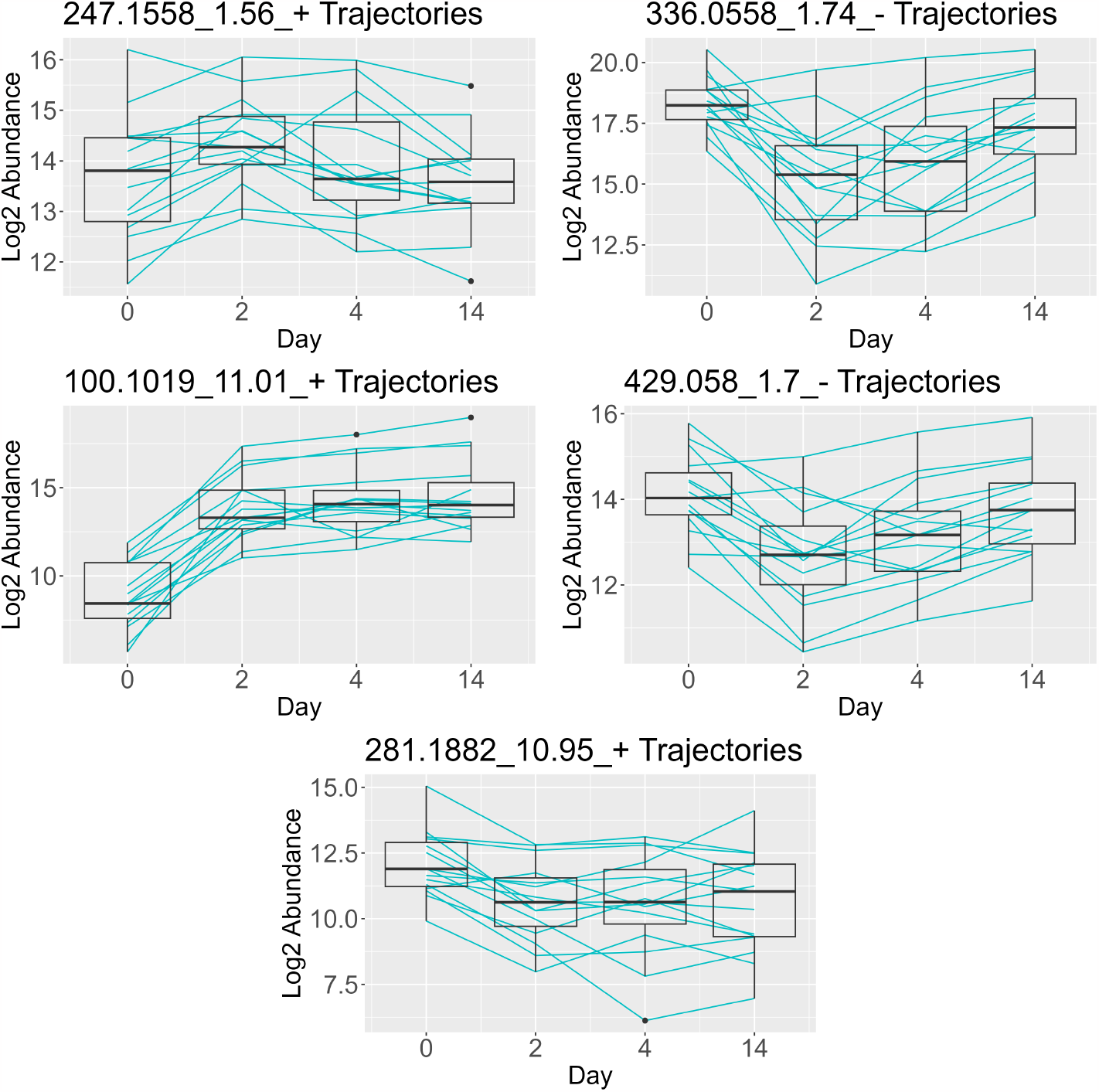
Trajectories for the 5 metabolites selected by PROLONG with the highest coefficients, excluding 204.1867 9.3 + shown in the introduction. These metabolites show a variety in trend and variance over time.

The univariate Wald tests select 116 of the metabolites, including 36 of the 45 selected by PROLONG. These additional selections may motivate further exploration of these metabolites, though the large-scale simulation results and the strength of the correlation among our metabolites suggest leaning towards the PROLONG results over the Wald tests.

## 5 Discussion

Data and figures shown in this paper illustrate the effectiveness of PROLONG as well as its univariate Wald test analog to a slightly lesser extent. PROLONG can be thought of as a principled classification method where we are not so much interested in the coefficients themselves or least squares error as we are in statistical power. The simulations show that both sensitivity and specificity are very high, particularly in the larger simulated datasets, and the real metabolites selected by PROLONG show the type of biological signal that motivated the model itself. Notably, PROLONG does not only select those metabolites that are strongly differentially expressed before and after treatment but also those that are relatively consistent in mean but show subject variability patterns that correspond to subject variability in improvement in TTP over time.

There is limited preprocessing necessary for PROLONG; all that is necessary is first-differencing the data, scaling the predictors, and computing the incidence matrix. Prior biological knowledge is helpful, and could be incorporated into the network penalty in place of absolute correlation in appropriate scenarios. By default, however, the network penalty uses observed correlations rather than known pathways and can be used as-is for data without a known connective structure or for data where molecule identification is possible but costly.

The extension of PROLONG to multiple types of continuous omics data is immediate. The graph and model assumptions do not require a specific covariance structure, so we should just combine the multiple omics types column-wise before proceeding with the same design matrix stacking and model procedure. For the graph edge weights, the correlation between two individual omics of different types is already interpretable, so we don’t expect to need any adjustments for this part of the extension. For a mixture of binary, count, and/or continuous omic data types, caution will need to be exercised in the selection of correlation measure for the graph’s edge weights. With an appropriate choice, the group lasso portion of the model should again fit in without any issues. This method extension should allow for effective multi-omic integration, not only allowing for simultaneous feature selection but also incorporating additional knowledge about how the features co-vary. The standard group lasso penalty used in PROLONG guarantees all nonzero or all zero coefficients within a group. There may be some advantages to inducing sparsity within each of these groups with sparse group lasso. In many datasets we should expect some time points, such as those closest to treatment, will be more important than others. Within-group sparsity may reveal additional information about the temporal relationship between the outcome and selected variables in such data.

The inclusion of continuous demographic variables should immediately work like with continuous multi-omics, and categorical demographic data should also work with some care in the selection of correlation measure. The design matrix Δ*X* would be the same as above but with a new demographic matrix *Z* appended after Δ*X*_2_. This way, the demographic data and its correlations will be used for each Δ*Y* exactly once. The network and lasso constraints would then apply in the same way, with each demographic variable forming its own group unless a specific grouping is desired.

If multi-omic integration is successful, the next obvious step would be expanding from strictly continuous response variables to count data with a Poisson or negative binomial GLM.

